# Single particle diffusion characterization by deep learning

**DOI:** 10.1101/588533

**Authors:** Naor Granik, Elias Nehme, Lucien E. Weiss, Maayan Levin, Michael Chein, Eran Perlson, Yael Roichman, Yoav Shechtman

**Affiliations:** Department of Biomedical Engineering, Technion - Israel Institute of Technology, Haifa 3200003, Israel; Lorry Lokey Interdisciplinary Center for Life Sciences and Engineering, Technion - Israel Institute of Technology, Haifa 3200003, Israel; Department of Electrical Engineering, Technion - Israel Institute of Technology, Haifa 3200003, Israel; Raymond & Beverly Sackler School of Chemistry, Tel Aviv University, Tel Aviv 6997801, Israel; Department of Physiology and Pharmacology, Sackler Faculty of Medicine, and Sagol School of Neuroscience, Tel Aviv University; Raymond & Beverly Sackler School of Physics, Tel Aviv University, Tel Aviv 6997801, Israel

## Abstract

Diffusion plays a crucial role in many biological processes including signaling, cellular organization, transport mechanisms, and more. Direct observation of molecular movement by single-particle tracking experiments has contributed to a growing body of evidence that many cellular systems do not exhibit classical Brownian motion, but rather anomalous diffusion. Despite this evidence, characterization of the physical process underlying anomalous diffusion remains a challenging problem for several reasons. First, different physical processes can exist simultaneously in a system. Second, commonly used tools to distinguish between these processes are based on asymptotic behavior, which is inaccessible experimentally in most cases. Finally, an accurate analysis of the diffusion model requires the calculation of many observables, since different transport modes can result in the same diffusion power-law α, that is obtained from the commonly used mean-squared-displacement (MSD) in its various forms. The outstanding challenge in the field is to develop a method to extract an accurate assessment of the diffusion process using many short trajectories with a simple scheme that is applicable at the non-expert level.

Here, we use deep learning to infer the underlying process resulting in anomalous diffusion. We implement a neural network to classify single particle trajectories according to diffusion type – Brownian motion, fractional Brownian motion (FBM) and Continuous Time Random Walk (CTRW). We further use the net to estimate the Hurst exponent for FBM, and the diffusion coefficient for Brownian motion, demonstrating its applicability on simulated and experimental data. The networks outperform time averaged MSD analysis on simulated trajectories while requiring as few as 25 time-steps. Furthermore, when tested on experimental data, both network and ensemble MSD analysis converge to similar values, with the network requiring half the trajectories required for ensemble MSD. Finally, we use the nets to extract diffusion parameters from multiple extremely short trajectories (10 steps).

## Introduction

Single-particle tracking (SPT) is widely used to investigate the biophysical properties of cellular membranes and other materials for extracting kinetics and other information on nano-scale processes. In recent years, rapid advances in labelling and detector sensitivity have widened the applicability for SPT to new biological systems with improved temporal and spatial resolution (1–4). The key information gained from these obtained trajectories after analysis is a statistical model for the mode of motion and the parameters which shed light on the dominating elements of the environment governing motion (1, 5, 6).

An essential strength of SPT methods is that the list of particle positions acquired during a measurement contains temporal information. This feature can be exploited to identify transient periods of statistically similar motion within the same trajectory, including different diffusion states (7–9), changes in diffusion type, *e.g.* distinguishing between Brownian, confined and directed diffusion (10–12) and the associated kinetics of transitions and equilibrium probabilities. While identification of periods of diffusive processes gives some insight into an objects behavior, we would ideally identify a specific mathematical model that best describes the measured trajectory.

Problematically, due to the stochastic nature of diffusive processes, a single trajectory does not uniquely correspond to a type of diffusion, unless It is long enough to show asymptotic behavior, for example, Fractional Brownian Motion (FBM) and Continuous Time Random Walk (CTRW) can both produce highly similar motion characteristics (3, 5), but arise from two completely different sets of underlying physical mechanisms. Therefore, it is usually only possible to assign a likely underlying mechanism by examining very long trajectories or alternatively, multiple intermediately long trajectories. This analysis is most frequently based on the mean-squared-displacements (MSD) curves, which describe the average spatial distances, ⟨*r*^2^⟩, measured between increasingly long time lags, *Δt* or *τ*, in the trajectory (13). This analysis has the benefit of being simple to apply, and the scaling behavior as a function of time follows the form of a power law ⟨*r*^2^⟩ ∝ *τ*^*α*^. For normal diffusion (*i.e.*, pure Brownian motion), *α* = 1, while for anomalous diffusion, the MSD can be subdiffusive (*α* < 1), or superdiffusive (*α* > 1), (Fig. 1) (14, 15).

**FIGURE 1.**
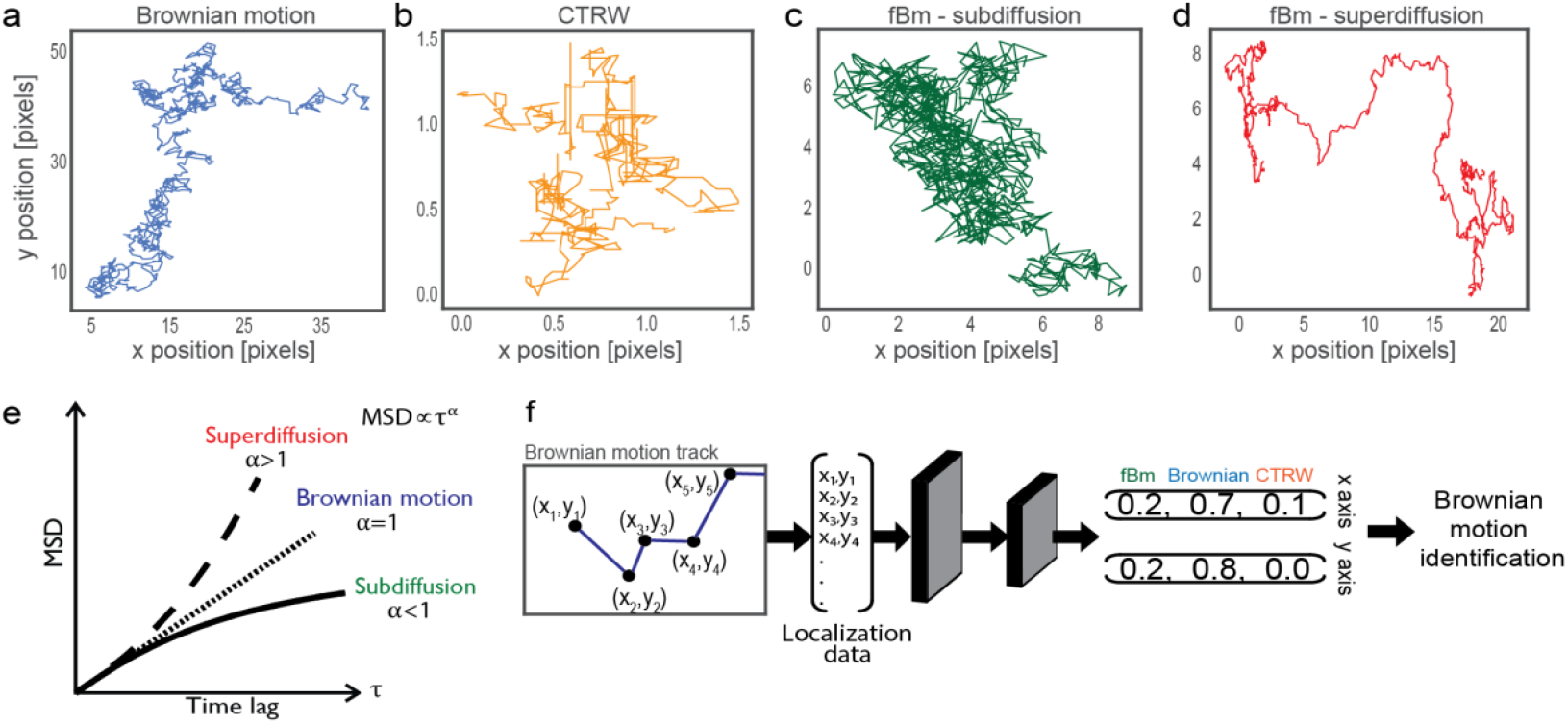
Diffusion models and analysis methods. **a-d**. Simulation examples of diffusion models discussed in this work. **e.** Mean Square Displacement curves for sub, super and normal diffusion. **f.** Schematic representation of the classification neural network.

Several typical families of processes leading to anomalous diffusion include fractional Brownian motion (FBM), Continuous Time Random Walk (CTRW), random walk on a fractal, and Lévy flights (3). Classifying a given trajectory to a diffusion model can reveal a great deal about the physical environment being investigated, for example, FBM might imply a crowded cellular environment, while CTRW can indicate an environment containing traps (16).

Existing methods for identification of the best-fitting diffusion model can be divided into two categories: qualitative, *e.g.* searching for ergodicity-breaking behavior or spatial constrains during motion; and quantitative, *e.g. p*-variation test, Gaussian-breaking parameters. Problematically, many of these criteria concern asymptotic behavior, and as such require a long single trajectory or hundreds of moderately long different trajectories from the same environment (16, 17).

Yet another issue is how to best determine specific-model parameters from a dataset for a given theoretical model. For example, using only the first two time lags of the MSD curve to determine the diffusion coefficient in pure Brownian trajectories was shown to be preferable under most conditions because it was more robust to noise (18). To the best of our knowledge, identifying the optimal approach for the relevant parameters describing subdiffusive motion has not yet been investigated, despite the known inaccuracies of applying traditional methods to deficient datasets (14, 15). Of particular interest is how to make use of short trajectories to accurately extract subdiffusive parameters, *i.e.* those produced by single-molecule fluorescence microscopy experiments using genetically-encoded labels. While these probes are seemingly ideal for reporting on the biological and biomimetic environments (19), their limited stability and brightness typically produces many short trajectories. The key challenge presented by these experiments is how to infer material properties of biological systems, and transport properties of single biomolecules from very short trajectories in which the asymptotic behavior is experimentally inaccessible.

The applicability of classical methods of accurately extracting the underlying parameters in this regime has been somewhat limited, thus necessitating a more reliable approach. Machine-learning algorithms, and in particular deep learning (20), excel at extracting concealed correlations in large datasets which can then be used to create a predictive tool for analysis of similar data (21). This makes the problem of single-particle diffusion characterization well suited for deep-learning analysis. Promisingly, several groups have applied classical machine learning (ML) (7) and deep learning (11, 12) algorithms to classify diffusion trajectories to confined, directed and normal diffusion, showing some advantage over traditional methods. For example, Muñoz-Gil and co-workers (22) recently addressed the challenge of anomalous diffusion characterization by using a Random Forest algorithm to classify a given trajectory as one of several anomalous diffusion models, and to estimate the anomalous diffusion exponent, as part of a classification problem, with a resolution of 0.1, achieving 70-90% accuracy depending to trajectory length and noise.

Here, we develop a deep-learning-based characterization framework for classification of diffusion processes in long trajectories, for which it exhibits higher precision over conventional analysis methods, as well as short and noisy trajectories, including parameter estimation from an ensemble of very short trajectories (10 time-steps). Our framework is comprised of multiple convolutional neural networks for classifying either single trajectories or a set of short trajectories as one of three selected diffusion models: Brownian motion, FBM and CTRW, and estimation of the relevant diffusion parameters by continuous regression. Our method is simple to implement and outperforms conventional approaches in terms of parameter estimation precision, convergence rate and usability of short trajectories.

## Results

For classification, we focus on three diffusion models which are commonly encountered in SPT experiments: Brownian motion, FBM and CTRW. We simulated each of the three models in order to produce the large amounts of data normally required to train a neural network.

Brownian motion was generated as a random walk process with independent identically distributed (IID) Gaussian steps (eq 1)

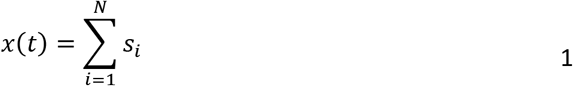

with {*s*_*i*_, *i* ≥ 1} a zero-mean Gaussian process.

Fractional Brownian motion is defined by its covariance function (eq 2)

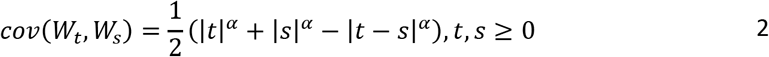

with {*W*_*t*_, *t* > 0}, a continuous zero-mean Gaussian process. Simulated tracks were generated using the circulant embedding algorithm (23).

Continuous time random walk can be regarded as a combination of random walks in both time and space, with temporal ‘steps’ (waiting times) drawn from a heavy-tailed distribution with an asymptotic power-law behavior (eq. 3).

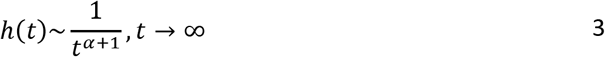

This general formulation has led to many variations over the years (24), we chose to implement the uncoupled CTRW simulation presented by Fulger et al (25), due to its relative simplicity and speed. In short, spatial increments are drawn from a symmetric *α*-stable levy distribution, temporal increments are drawn from a Mittag-Leffler distribution.

For experimental validation of our methods, we performed three experiments, each demonstrating a different diffusion model: Diffusion of fluorescent beads in Actin networks, demonstrating FBM, with a different H parameter as a function of Gel mesh density (26, 27); Diffusion of fluorescent beads in a water-Glycerol solution, demonstrating Brownian motion; Diffusion of proteins on membrane surfaces demonstrating FBM subordinated to CTRW (28).

We trained a neural network to classify a given single particle trajectory to one of the three diffusion models described above. The network was trained on ∽300,000 simulated trajectories of 100 steps, with Gaussian localization error. For FBM and CTRW, which are parameter dependent, each simulated trajectory was generated with a random parameter drawn from a uniform distribution in the range [0,1]. Both models converge to Brownian motion for specific parameter values (0.5 for FBM and 1 for CTRW); to account for this, trajectories generated with a parameter in the range [0.4,0.6] for FBM, and [0.9,1] for CTRW, were labeled as Brownian motion.

During testing, the network receives 2D tracks as two separate 1D vectors (x and y data) and outputs the probabilities of being associated to each of the above diffusion models. This configuration has the advantage of axial separation – in general, the diffusion along the different axes is uncorrelated – however, a diffusing particle presenting different motion characteristics in separate directions (due to environmental constraints for example) will be easily detected by the network. The performance of the trained network was tested on simulated data with various levels of Gaussian noise and with Ornstein-Uhlenbeck noise which is used as a model for “environmental noise” (see definition in Supporting Material) (25).

The classification net achieves excellent performance on simulated data. As seen in figure 2a, it is able to correctly identify FBM and CTRW tracks, in various SNR levels (Fig. S2), and parameter selection (disregarding parameter ranges *H* ∈ [0.4,0.6] and *α* ∈ [0.9,1] which are equivalent to Brownian motion). The network also correctly identifies experimental data, as seen in Fig. 2b, where three sample tracks from different experiments are correctly characterized to different diffusion models - FBM for bead in a crowded actin gel (250 nm mesh size); Brownian motion for bead in 40% glycerol solution; and FBM combined with CTRW for transmembrane protein diffusion. Additional results can be found in the supplementary material.

**FIGURE 2.**
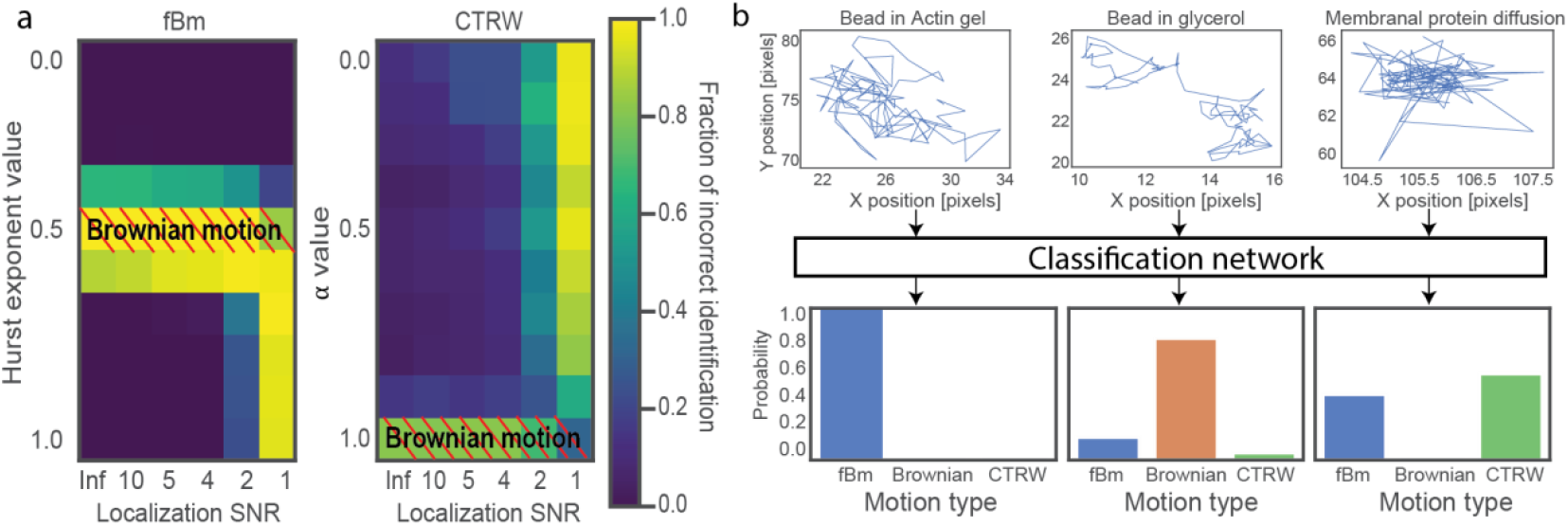
Model identification (classification) network. **a.** Heat maps presenting fraction of classification errors as a function of model parameter and SNR. Each pixel represents average results from 200 simulated trajectories. **b.** Network analysis for three different sample tracks – from left to right: Fluorescent bead diffusing in actin gel demonstrating FBM; fluorescent bead diffusing in glycerol solution, demonstrating Brownian motion; Membranal protein diffusion presenting a combination of FBM and CTRW.

For estimation of the Hurst exponent (H), the parameter of interest in FBM (29), we trained a set of neural networks on ∽150,000 simulated tracks with randomly selected Hurst exponents in the range [0.05,0.95]. Two separate versions were trained: first, an array of single-track networks (ST-networks), which receive as input the autocorrelation of the derivative of a single 1D trajectory, known as the velocity autocorrelation. Each network in the set was optimized for different track lengths: from 25 to 1000 steps. The networks were tested on simulated data with various levels of added noise and compared to time averaged MSD (TAMSD) estimation. On simulated data, the ST-networks proved to be superior to TAMSD in terms of both estimation accuracy, the number of steps required, and robustness to noise. This is most evident in Fig. 3a, b, which presents a comparison between the network optimized for 100 step-tracks and TAMSD. The results show a bias in TAMSD estimation (Fig. 3b – blue line underestimates true Hurst exponent), which can result from the relatively short track lengths. In comparison, the network is able to correctly estimate the Hurst exponent, and with lower standard deviation; further results can be found in supplementary material.

**FIGURE 3.**
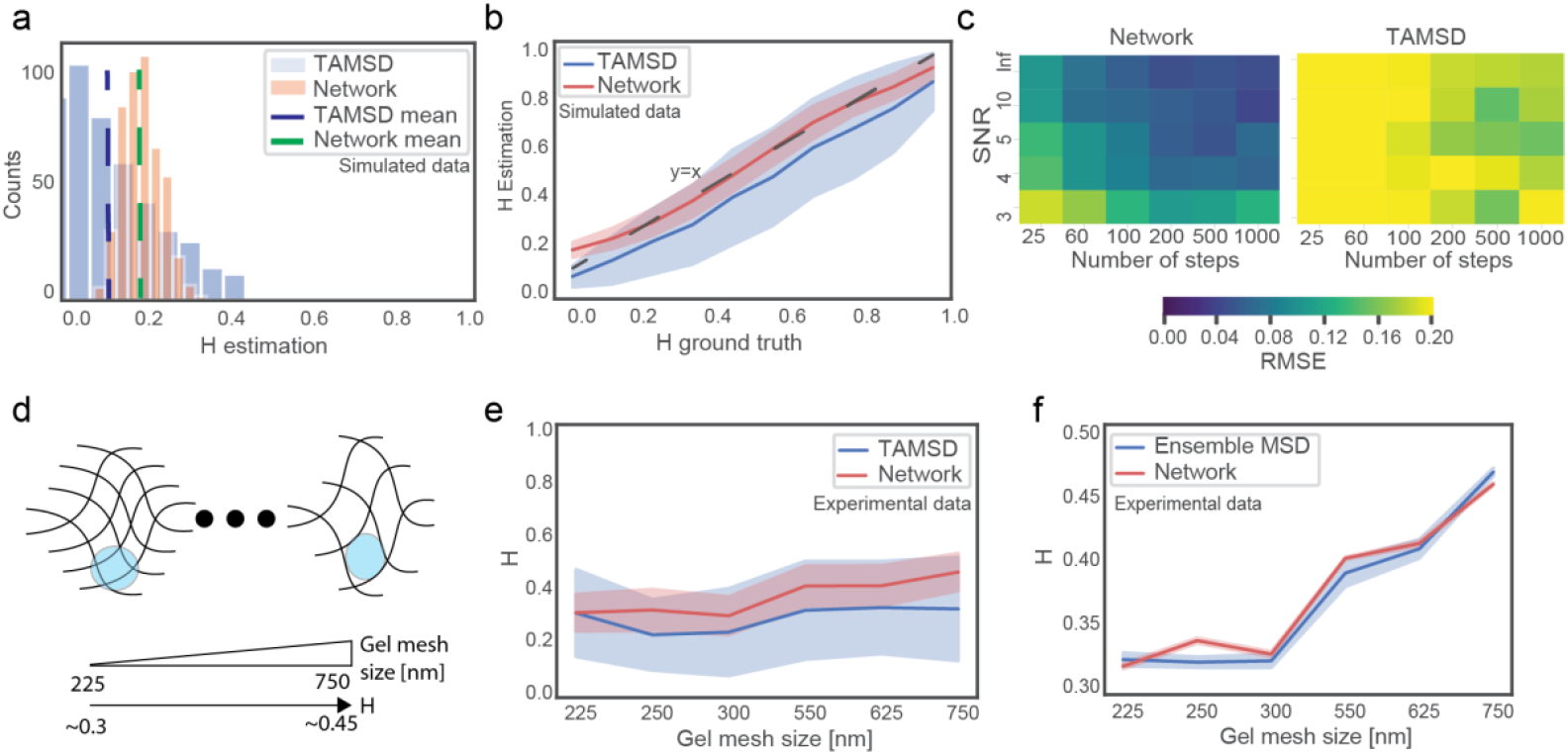
Hurst exponent (H) estimation network. **a**. Comparison to time averaged MSD - Estimation of single value, 200 tracks were simulated with the same Hurst exponent (0.2). Estimation was done per track by both 100-step network and TAMSD. **b.** Comparison to time averaged MSD - Estimation on range, for each H in range [0.05,0.15,..0.95], 1000 tracks were generated with SNR = 4, and estimated by 100-step network and TAMSD. Results presented are estimation mean and standard deviation. **c.** RMSE heat maps between estimation to ground truth for simulated tracks as a function of SNR and number of steps in a track. Each pixel represents 100 tracks with random H values in the range [0,1]. **d.** Schematic of experimental data. Depicted are beads “stuck” in actin networks, ratio of bead size to gel mesh size determines the value of Hurst exponent. **e.** Performance on experimental data vs. time averaged MSD, estimation based on 100 steps only. **f.** Performance on experimental data vs. ensemble MSD, estimation of each experiment is performed by the network most fitting the length of the experimental tracks (typically thousands of steps per trajectory).

Much like in MSD, estimation is better when more data is available, i.e. additional steps (Fig. 3c). Thus, when a 500-step trajectory is available, the networks optimized to run on longer tracks (100, 200, or 500) will outperform a network optimized for 25 steps.

Performance of the H-estimation network was tested on experimental data sets of fluorescent beads diffusing in entangled F-actin networks gel with various mesh sizes (see (27) for details of preparation), allowing control of the Hurst exponent by changing the crowding of the environment (Fig. 3d).

In this case, due to the lack of a ground truth, the net’s estimation was compared to that of TAMSD and ensemble MSD. Compared to TAMSD analysis (Fig. 3e), the network estimates values slightly higher than TAMSD, with the exception of the experiment performed in 225 nm mesh size. For all data points, network estimation is well within the TAMSD standard deviation (STD), with its own STD being less than half that of MSD (∽0.06 vs 0.15). Compared to ensemble MSD (Fig. 3f), network and MSD estimations converge to relatively equal values (within ±0.01), with the network exhibiting lower standard deviation (0.001 vs 0.005). (See Fig. S6 for complete results).

The second version of the H-estimation network aims to tackle a problem of high practical importance of experiments in which numerous very short trajectories are available (i.e. ∽10 steps), rather than a single long trajectory. This is often the case when tracking fluorescent proteins that are quick to photobleach (30). To this end we trained an array of multi-track networks (MT-networks), that receive as input matrices of 1D-velocity autocorrelations of 10-step trajectories (Fig. 4a). The MT-networks were compared to ensemble MSD estimation on simulated data and proved to outperform it in terms of accuracy, standard deviation and convergence speed (Fig. 4, b and c). It is important to note however, that when allowing trajectories of longer lengths, ensemble MSD ultimately surpasses network performance in terms of accuracy.

**FIGURE 4.**
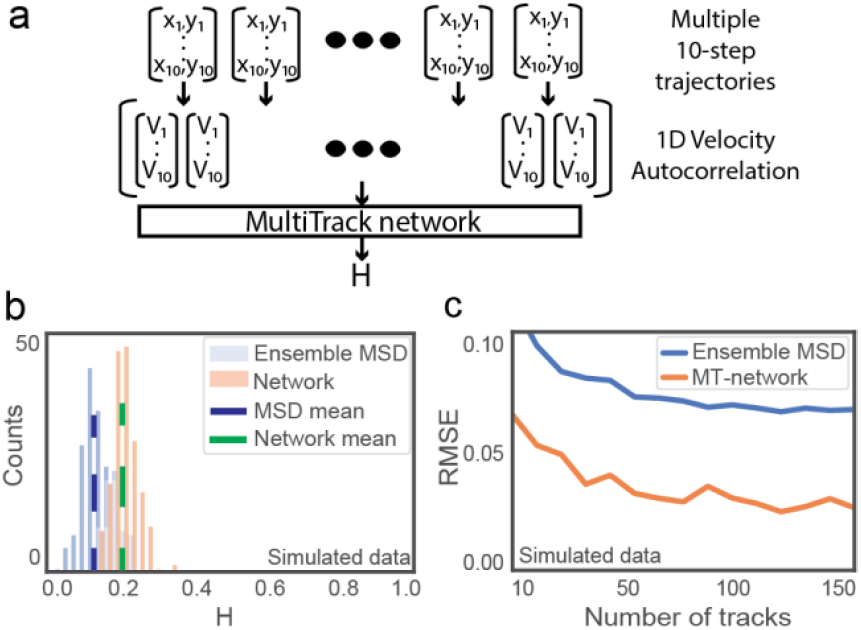
Multi-track networks. **a**. Schematic of MT-network **b.** Estimation of single value. 200 matrices of 50×10 tracks were generated with H=0.2 and analyzed by ensemble MSD and network. **c.** RMSE estimation to ground truth, for each value M in x-axis, 500 matrices of Mx10 tracks were generated with H=0.2 and analyzed by ensemble MSD and MT-networks.

For the problem of diffusion coefficient estimation from Brownian motion trajectories, we trained a single neural network which receives as input two scalar values – the mean of the absolute velocity, and its standard deviation. We found this combination holds powerful predictive capabilities for short trajectories. We note, that for long trajectories these two observables are the same.

Training was performed on ∽100,000 tracks of 1000 steps, with diffusion coefficients randomly drawn from a uniform distribution in the range [0.1,10] 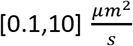. For a single value, the network was found to have a lower standard deviation than TAMSD on tracks of only 50 steps (Fig. 5a). When considering the range of possible values, the network was discovered to be precise on the low to medium range (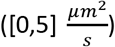) as seen in figure 5b where estimation results converge with those of TAMSD. For this problem, no MT-networks were trained; Multi-track analysis is unnecessary in this case, due to the fact that Brownian motion is a memoryless process, and therefore different tracks from the same population can be concatenated and analyzed in the same manner. Using this concatenation method, the network was compared to ensemble MSD estimation on collections of 10-steps tracks and was found to be more accurate regardless to the number of tracks (Fig. 5c).

**FIGURE 5.**
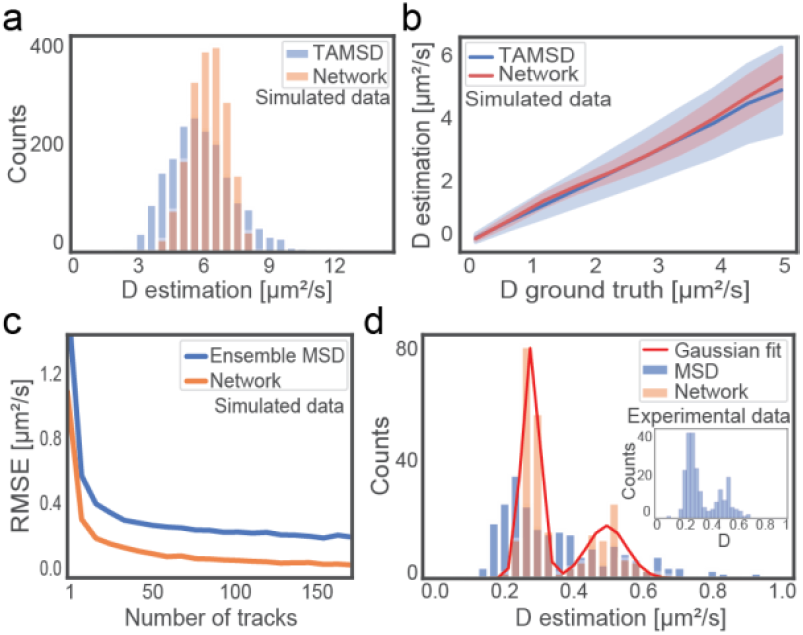
Diffusion coefficient estimation network. **a**. Estimation of single value. 2000 tracks of 1000 steps were simulated with the same D values (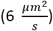). Estimation was done per track by both network and time average MSD based on 50 steps only. **b.** Performance on a range of values. For each value in x-axis, 1000 tracks of 100 steps were simulated with the corresponding D-value, presented are the mean and STD of estimations by both time averaged MSD and network **c.** Comparison to ensemble MSD on very short trajectories. For each value M in x-axis, 500 matrices of Mx10-step-tracks were generated with D=3. Performance measured as error from ground truth. **d.** Experimental data. Two populations of beads diffusing in glycerol solution. Estimation results for network and time averaged MSD are based on 100 step segments of the full tracks. Inset: Time averaged MSD estimation on the full 600 step-tracks.

Performance was tested on experimental data of fluorescent beads of two sizes (100 nm and 200 nm) diffusing in 40% glycerol solution (see (31) for details of preparation). The results, based on 100-step tracks, are presented in figure 5d. Network estimation shows two different populations, with mean values of 0.29 and 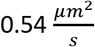, which are similar to predicted theoretical diffusion coefficients calculated from the Stokes-Einstein equation – 0.27 and 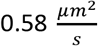. Time-averaged MSD estimation of the same data shows relatively close values; however, it cannot correctly separate the two existing populations (see figure 5d inset for TAMSD estimation of the full tracks – using 500 step long trajectories)

## Discussion

Deep learning is revolutionizing signal analysis in numerous scientific fields, owing to its ability to identify complex models in large quantities of data, relative simplicity, (once trained) inference speed, and robustness. This revolution is providing tools enabling the extrapolation of biological conclusions from seemingly unintelligible measurements. In this work, we take a first step in strengthening single particle diffusion analysis using a set of neural networks for model selection and parameter estimations. We have shown these to be more precise than current standard methods on both simulated and experimental data, while requiring a smaller number of steps, with increased robustness to noise and the advantage of being parameter free.

The framework we provide allows for almost trivial concatenation of different neural nets, to provide end-to-end classification followed by diffusion parameter estimation. Future work in this field should expand the set of networks to include other models, e.g. estimation of CTRW parameters, identification of motion on fractal, and levy flights, and to address cases of a hierarchy of transport modes manifested in the same trajectory.

## Supporting citations

References (32–38) appear in the Supporting Material

## Author Contributions

N.G. performed research and analysis and wrote the paper. Experiments were performed under the supervision of E.P, Y.R and Y.S. By M.L, M.C and L.E.W. Y.R. and Y.S. advised and supervised this work and wrote the paper.

## Acknowledgements

This research was supported by the Israel Science Foundation (grant No. 450/18). LEW and YS are supported by the Zuckerman STEM Leadership Program, and YS by a Career Advancement Chair from the Technion’s Leaders in Science and Technology program.

