## Supplementary Material for "Single particle diffusion characterization by deep learning" (from another version of this preprint: version 1, 2019-03-26)

**Table of contents**

1. CNN network architecture
2. SNR definition
3. Classification network – additional results
    a. Confusion matrices
    b. Classification performance with Ornstein-Uhlenbeck noise
    c. Classification of experimental data – beads in glycerol solution
    d. Classification of experimental data – Protein diffusion
4. H regression network – additional results
5. Multi-Track network – additional results
6. D regression network – data conversion
7. Beads in glycerol experiments
    a. Theoretical calculation
    b. Movie S1
8. Materials and methods
    a. Fluorescent beads in F-actin networks
    b. Fluorescent beads in Glycerol solution

### 1. CNN network architecture

Network architecture is based on the design shown in (1). Four sets of convolution blocks with different filter sizes [2,3,4,10], operate in parallel (Fig S1A). Each block consists of 1D dilated causal convolution layers with increasing dilation factors (Fig S1B). This setup was designed to find correlations spanning multiple time scales of unknown length. This architecture was selected in a process of trial and error, based on classification and regression performance on simulated data. Additional convolution layers were added or removed according to the track size a specific network was intended for. For example, the network designed for 1000-step tracks has an additional convolution block with filter size of 20.

For the Multi-Track networks, the 1D convolution layers were replaced by 2D convolution layers with dilation factors operating on the temporal axis only (i.e. for an input matrix M with the shape [Number of tracks][Number of steps] the dilation factor will be (1,d)).

Networks were implemented and trained using Keras (Version 2.2.4) with TensorFlow backend version (1.8.0) in Python (version 3.5). Other packages used: NumPy (version 1.14.5), SciPy (version 1.2.1), Stochastic (version 0.4). Training was done on NVIDIA GeForce Titan GTX in a Windows environment.

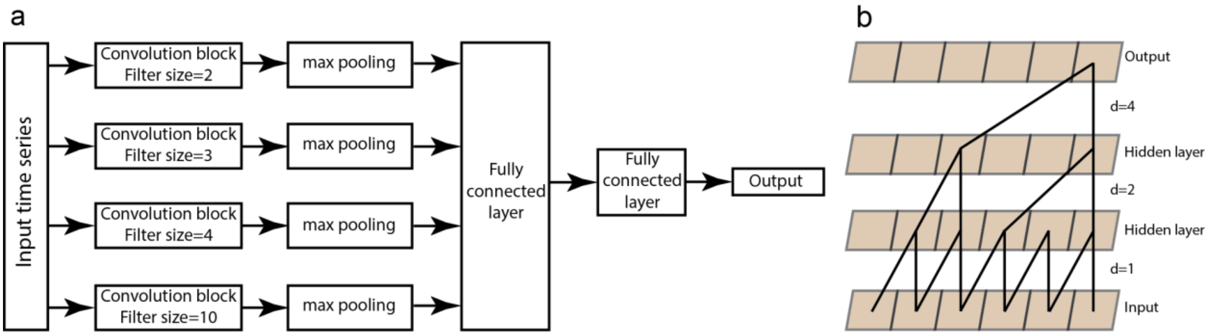


FIGURE S1. Neural network architecture. **A.** Schematic of the neural network basic structure. **B.** An example convolution block with filter size = 2 and dilation factors $d = 1,2,4$.

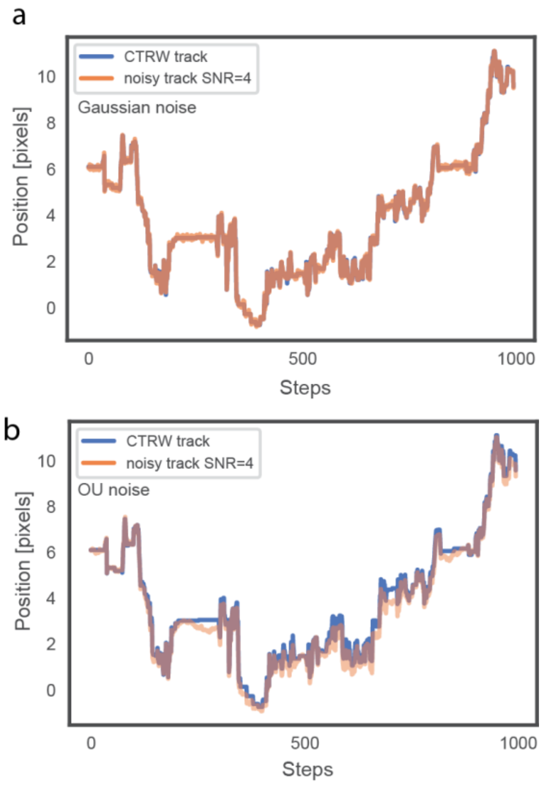


FIGURE S2. Sample noisy tracks. **a.** CTRW track (blue) with added Gaussian noise(orange), SNR = 4. **b**. CTRW track (blue) with added OU noise (orange). SNR = 4

### 2. SNR definitions

Localization noise: For a trajectory $x$, we define the Signal to Noise Ratio (SNR) as the ratio between the standard deviation of the signal increments to the standard deviation of the Gaussian noise added to the signal.

$$SNR \triangleq \frac{std\left(\frac{dx}{dt}\right)}{std(N)}$$

For $N$, a zero mean Gaussian process.

A second noise process which we consider is the Ornstein-Uhlenbeck (OU) noise, which can reflect environmental noise (i.e. an active environent “pushing” against the diffusing particle) (2, 3).

In simple terms, an OU process can be considered as a Brownian motion with an additional feedback relaxation to a mean position $\mu$. Mathematically, $U_t$ is an OU process if it satisfies the following stochastic differential equation:

$$dU_t = \theta U_t(\mu - t)dt + \sigma dW_t$$

With $W_t$ a zero mean Gaussian process. $\theta$ – speed of relaxation; $\mu$ – mean of the process; $\sigma$ – volatility of the process.

We consider an OU process with $\mu = 0, \theta = \sigma = 1$. For a trajectory $x$, we define the noisy trajectory as:

$$x_n = x + kU_t, \;\; k \geq 0$$

and the SNR as:

$$SNR = \frac{1}{k}$$

### 3. Classification network - additional results

#### a. Confusion matrices

Classification confusion tables presented are organized according to SNR levels and are all based on simulated tracks of 100 steps. Tables were produced by simulating a set of 300 tracks, 100 for each diffusion model. Parameters for CTRW and FBM were selected at random from the range of values that should not result in Brownian motion ($\alpha \in [0.05, 0.9], H \in [0.05, 0.45] \cup [0.55, 0.95]$), in order to maintain correct statistics in the data set.

| **SNR = ∞** | | Ground truth | | |
|---|---|---|---|---|
| | | FBM | Brownian | CTRW |
| Network prediction | FBM | 84 | 14 | 2 |
| | Brownian | 0 | 99 | 1 |
| | CTRW | 5 | 2 | 93 |

| **SNR = 10** | | Ground truth | | |
|---|---|---|---|---|
| | | FBM | Brownian | CTRW |
| Network prediction | FBM | 82 | 16 | 2 |
| | Brownian | 0 | 99 | 1 |
| | CTRW | 6 | 3 | 91 |

| **SNR = 5** | | Ground truth | | |
|---|---|---|---|---|
| | | FBM | Brownian | CTRW |
| Network prediction | FBM | 80 | 17 | 3 |
| | Brownian | 0 | 99 | 1 |
| | CTRW | 9 | 5 | 86 |

| **SNR = 2** | | Ground truth | | |
|---|---|---|---|---|
| | | FBM | Brownian | CTRW |
| Network prediction | FBM | 77 | 20 | 3 |
| | Brownian | 26 | 74 | 0 |
| | CTRW | 28 | 12 | 60 |

| **SNR = 1** | | Ground truth | | |
|---|---|---|---|---|
| | | FBM | Brownian | CTRW |
| Network prediction | FBM | 67 | 31 | 2 |
| | Brownian | 99 | 1 | 0 |
| | CTRW | 69 | 23 | 8 |

The confusion matrices show the identification network is accurate even at relatively low SNR levels, beginning to falter at SNR=2. Another important result is the uncertainty between FBM and Brownian motion, even with no addition noise. This is caused by the fact that FBM is a generalization of Brownian motion, with certain parameter choices causing the network to err between the two. This is true despite the fact that the Brownian motion parameter range – [0.45-0.55] was not used during generation of the dataset.

To illustrate this, we show below the confusion table for SNR=∞, but for a data set wherein H was selected from the parameter range – $[0.05,0.35] \cup [0.65,0.95]$.

| **SNR = ∞** **Reduced H range** | | Ground truth | | |
|---|---|---|---|---|
| | | FBM | Brownian | CTRW |
| Network prediction | FBM | 93 | 7 | 0 |
| | Brownian | 0 | 100 | 0 |
| | CTRW | 8 | 1 | 91 |

An additional result arising from the above tables, is that there is no noticeable ambiguity between CTRW and Brownian motion. This possibly has less to do with parameter choices, but rather with the features the neural network learns. During the training phase, each filter learns different features of the signal, CTRW is characterized by long waiting periods between jumps, which results in the diffusing particle being noticeably 'stuck' in its position. It is highly likely that the network found this significant, resulting in signals in which there is some evidence of a stuck particle being classified as CTRW. A further indication to this can be found in the confusion table for SNR=1, in which the additional noise masks the signal itself, making it similar to FBM.

**b. Classification accuracy with Ornstein-Uhlenbeck noise**

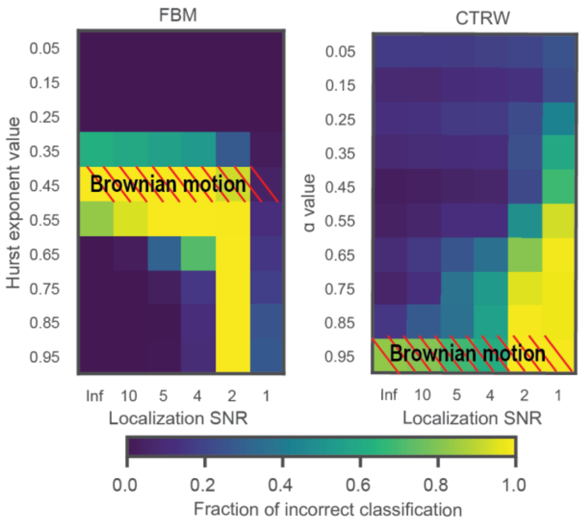


FIGURE S3: Model identification (classification) network with additional OU noise. Heat maps presenting fraction of classification errors as a function of model parameter and OU-SNR. Each pixel represents results from 200 simulated trajectories.

Figure S3 presents the fraction of wrong predictions as a function of parameter and SNR. For CTRW, the network performs well up to SNR of 4, where it begins to falter, reverting to FBM due to the additional noise process, however for low values of $\alpha$, the network retains its CTRW prediction.

For FBM, we see similar behavior to the case of Gaussian localization noise, with the exception of SNR = 1, where we see evidence that the noise process is a fractional Gaussian process, comparable to FBM (3).

To further illustrate this, presented below are two confusion matrices for SNR = 5,1.

| **SNR = 5** | | Ground truth | | |
|---|---|---|---|---|
| | | FBM | Brownian | CTRW |
| Network prediction | FBM | 86 | 14 | 0 |
| | Brownian | 0 | 99 | 1 |
| | CTRW | 9 | 5 | 86 |

| SNR = 1 | | Ground truth | | |
|---|---|---|---|---|
| | | FBM | Brownian | CTRW |
| Network prediction | FBM | 91 | 7 | 2 |
| | Brownian | 97 | 3 | 0 |
| | CTRW | 62 | 7 | 31 |

### c. Classification of experimental data set – beads diffusing in glycerol solution

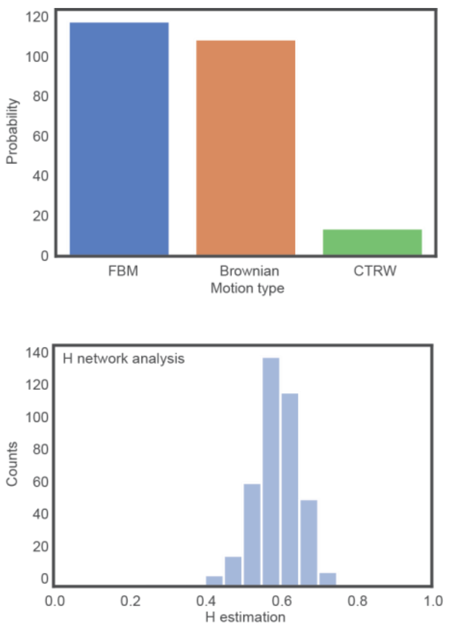


FIGURE S4: Classification of experimental data – beads diffusing in glycerol. Top: Classification results. Bottom: analysis of same data by H-network based on 100 steps.

Figure S4 shows classification results for the experimental bead-in-glycerol data. As can be seen, the classification is not perfect, showing nearly similar numbers of FBM and Brownian motion (the minor CTRW population represents beads stuck to the surface unable to move, these do not appear in the H-estimation analysis). The fault most likely lies in a combination of precision errors and other unknown factors relating to the experiment (e.g. effects of fluid dynamics). Analysis using H network shows a population centered around H = 0.6 with standard deviation of 0.07, in agreement with the classification results (i.e. approximately half classified as FBM, and half as Brownian motion).

### d. Classification of experimental data set – Proteins diffusing on a membrane surface

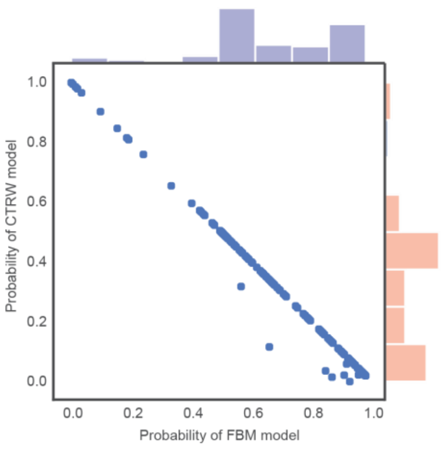


Figure S5: Classification of experimental data – Proteins diffusing on membrane surface. Results presented are probabilities of being identified as FBM model (x axis) or CTRW model (y axis)

This experiment presents a unique challenge in that the motion does not fit into any one anomalous diffusion model. For this reason, we cannot simply set the highest probability in the network output as the selected model, but instead must look at probabilities themselves. Fig. S5 shows the 2D probability distribution space from 205 experimental trajectories. X, Y axes represent probabilities of being assigned to FBM and CTRW models, respectively. The data closely follows a $y = -x$ trend line, with clusters of tracks being scattered around (x,y) = (0.5,0.5), or (x,y) = (1,0) From this we can conclude: The network identifies features from both models, while almost entirely disregarding the Brownian motion model (otherwise the sum of $P_{fbm}$ and $P_{CTRW}$ would not be one); The network shows a bias towards FBM as was previously shown on simulated data.

### 4. Hurst parameter regression network - additional results

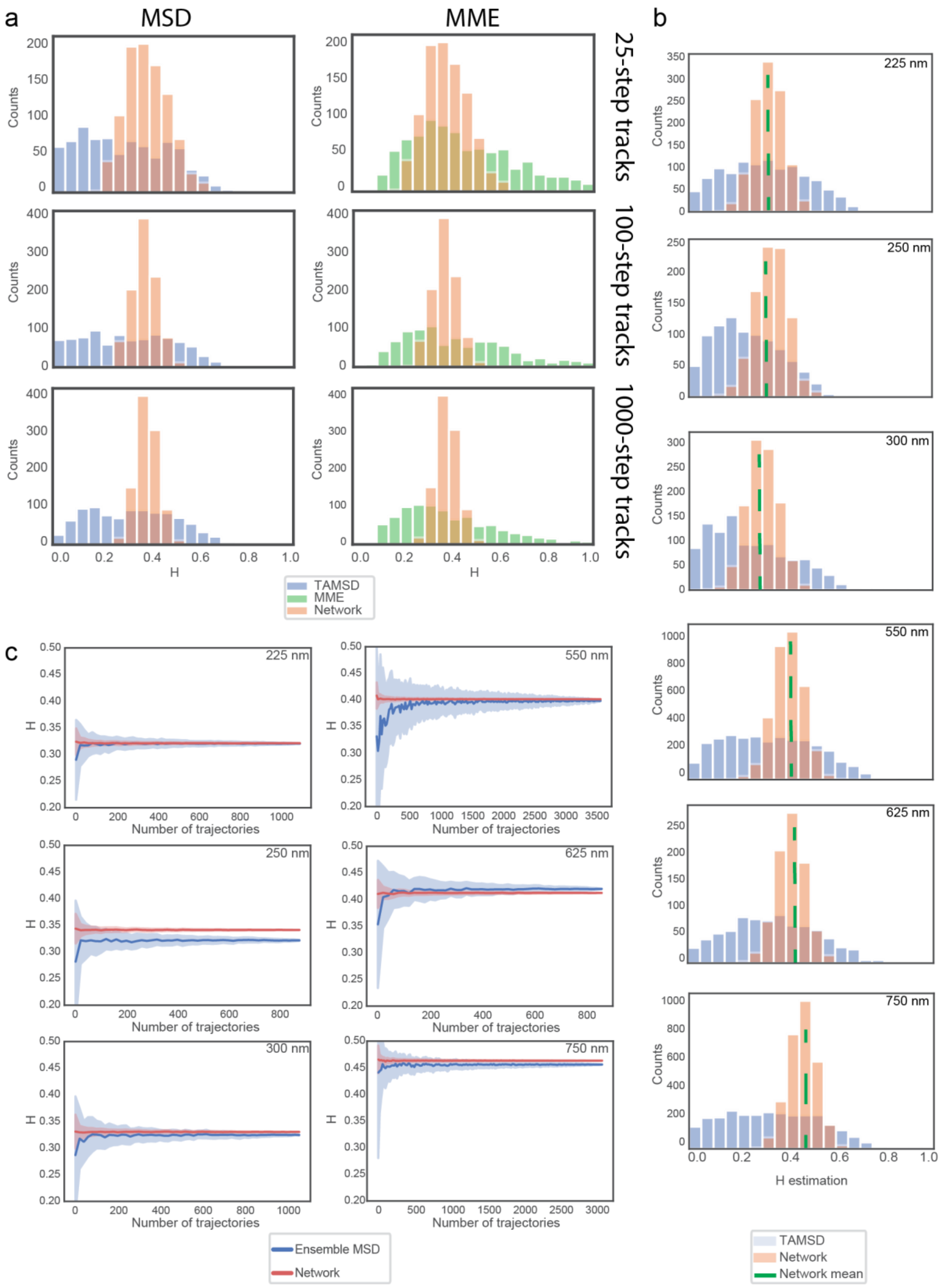


FIGURE S6: Additional H network results. **A.** Estimation of single value by TAMSD, MME and networks on tracks of different sizes. 1000 different tracks were generated with H = 0.4 and SNR = 4 **B+C.** Experimental data – beads diffusing in Actin gel of different mesh sizes (sizes written in figure). B – Evaluation by TAMSD and network; C – Evaluation by ensemble MSD and network.

Figure S6-A presents a comparison between three methods – Mean Square Displacement (MSD), Mean Maximal Excursion (MME) (4) and network estimation for three different track lengths. Tracks were simulated with H = 0.4 and SNR = 4. Estimation is based on single tracks only. The results are summarized in the following table:

| | 25-steps | 100-steps | 1000-steps |
|---|---|---|---|
| Network | $0.39 \pm 0.08$ | $0.37 \pm 0.05$ | $0.39 \pm 0.04$ |
| MSD | $0.22 \pm 0.21$ | $0.29 \pm 0.19$ | $0.32 \pm 0.17$ |
| MME | $0.55 \pm 0.41$ | $0.48 \pm 0.31$ | $0.45 \pm 0.3$ |

The results show that all three methods converge to an estimation of 0.4, as the tracks increase in length, with the network outperforming MSD and MME in both accuracy and estimation standard deviation.

Figure S5-B, C present the complete results for the experimental data summarized in the paper. Comparing network estimation to time averaged MSD and ensemble MSD, respectively.

### 5. Multi-Track network - additional results

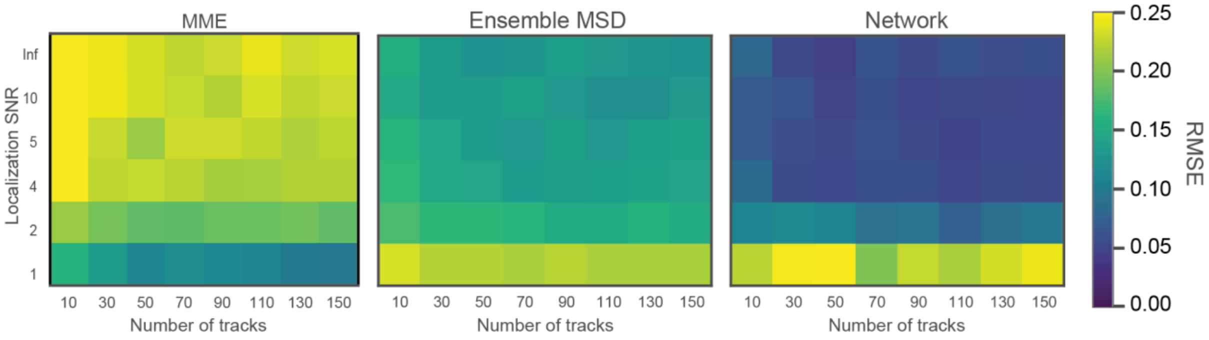


FIGURE S7: Multi-Track network heat maps showing RMSE as function of Localization SNR and number of tracks used for MME, ensemble MSD and MT-network. Each pixel represents 200 simulations of Mx10-step-tracks generated, with M being the number of tracks corresponding to the value in the heat map x-axis.

### 6. Diffusion coefficient regression network - data conversion

D-network was defined to operate at a specific pixel size and frame length, 550nm and 0.05 seconds, respectively. These settings were selected to fit available experimental data. Due to the nature of the problem, input data cannot be standardized as this would effectively destroy the diffusion coefficient information in the data. Despite this hard-coded setting, data from different setups can be entered with two simple conversion steps.

Temporal conversion can be done post-analysis, on the diffusion coefficient results by the equation:

$$D = D_{network} \cdot \frac{0.05}{\Delta t_{experiment}}$$

Pixel size conversion must be done on the localization data, prior to calculating the mean and standard deviation of increments, this is done by the equation:

$$x_{corrected} = x_{experimental} \cdot \frac{experimental\ pixel\ size}{0.55}$$

Following conversion, the data can be entered to the network with the output being in $\frac{\mu m^2}{s}$.

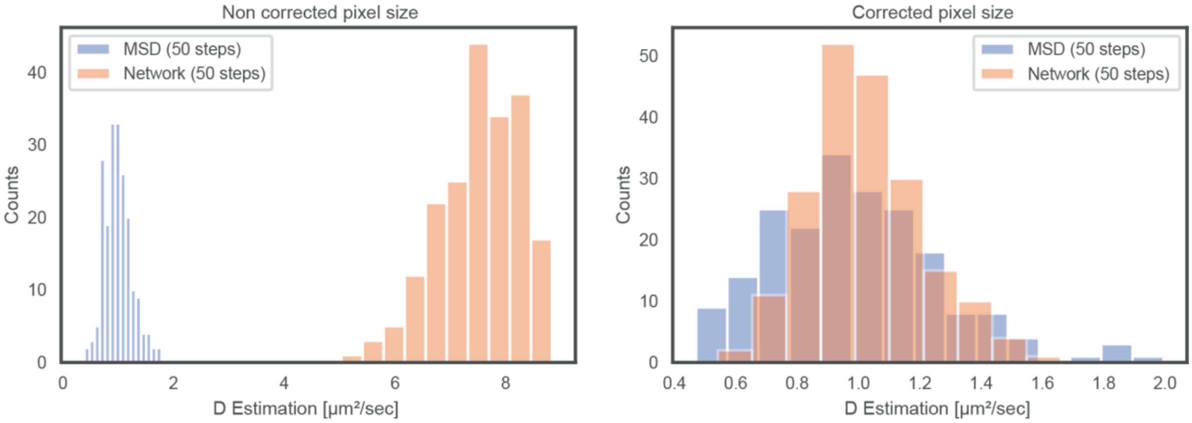


FIGURE S8: Effect of pixel conversion. 200 Tracks were generated with $D = 1[\frac{\mu m^2}{s}]$, with pixel size setting of 200 nm per pixel and analysed by the network and by temporal MSD. Left: Estimation of raw track data, without pixel size correction, **MSD was calculated with correct pixel size.** Right: Estimation of track data with pixel conversion.

### 7. Beads in glycerol experiment

#### a. Theoretical calculations

Theoretical diffusion coefficient values for the diffusion-in-glycerol experiment were calculated using the Stokes-Einstein equation for diffusion of spherical particles through a liquid with low Reynolds number (5).

$$D = \frac{K_B T}{6\pi\eta r}$$

Where:

$D$ – Diffusion coefficient $\left[\frac{\mu m^2}{s}\right]$

$K_B$ – Boltzmann constant $\left[\frac{m^2 kg}{k \cdot s^2}\right]$

$T$ – Temperature $[k]$

$\eta$ – Viscosity $\left[\frac{kg}{m \cdot s}\right]$

$r$ – Particle radius $[nm]$

The experiments were conducted in room temperature with beads of two different sizes - 100,200 $[nm]$ in a solution of 40% glycerol in water, giving a viscosity coefficient of 0.00372 $\left[\frac{kg}{m \cdot s}\right]$ (6).

$$D_{100[nm]} = \frac{K_B T}{6\pi\eta r} = \frac{1.3806 \cdot 10^{-23} \cdot 293}{6\pi \cdot 0.00372 \cdot 100 \cdot 10^{-9}} = 5.722 \cdot 10^{-13} \left[\frac{m^2}{s}\right] = 0.57 \left[\frac{\mu m^2}{s}\right]$$

$$D_{200[nm]} = \frac{K_B T}{6\pi\eta r} = \frac{1.3806 \cdot 10^{-23} \cdot 293}{6\pi \cdot 0.00372 \cdot 200 \cdot 10^{-9}} = 2.884 \cdot 10^{-13} \left[\frac{m^2}{s}\right] = 0.28 \left[\frac{\mu m^2}{s}\right]$$

**b. Supporting movie**

Movie S1 depicts a sample experiment consisting of two populations of beads diffusing in 40% Glycerol solution. Green and red boxes mark beads with radiuses of 100 and 200 nm respectively.

### 8. Materials and methods

**a. Fluorescent beads in F-actin networks**

We prepare F-actin networks as was described previously (7, 8). We determine the mesh size from the concentration of the actin monomer according to $\xi_s = 0.3\sqrt{C_A}$ (9).

We use a=0.55 µm polystyrene beads (Invitrogen Lot \#742530) and no capping protein.

**b. Fluorescent beads in Glycerol solution**

Freely-diffusing, bead-tracking experiments were performed as described previously (10). In brief, a passivated diffusion chamber was prepared by treating the surface of a glass slide and coverslip with 20 mg/mL casein solution in PBS. The passidvation solution was then removed and replaced with diluted fluorescent-microsphere (100 and 200 nm diameter fluorospheres, Life Technology) diluted in 40% glycerol in water (v/v). The chamber was then sealed with nail polish, then imaged using a standard inverted microscope system (TI Eclipse, Nikon) with a 20X objective (DETAILS, Nikon) using an sCMOS detector (Photometrics). Movies were recorded with 50 ms frames using NIS Elements software (Nikon) and analyzed using XYZ.
